# Adaptive coding of pain prediction error in the anterior insula

**DOI:** 10.1101/2021.10.15.464508

**Authors:** R Hoskin, D Talmi

## Abstract

**Background:** Understanding the mechanisms behind the influence of expectation and context on pain perception is crucial for improving analgesic treatments. Prediction error (PE) signals how much a noxious stimulus deviates from expectation and is therefore crucial for our understanding of pain perception. It is thought that the brain engages in ‘adaptive coding’ of pain PE, such that sensitivity to unexpected outcomes is modulated by contextual information. While there is behavioural evidence that pain is coded adaptively, and evidence that reward PE signals are coded adaptively, controversy remains regarding the underlying neural mechanism of adaptively-coded pain PEs.

**Methods:** A cued-pain task was performed by 19 healthy adults while undergoing FMRI scanning. BOLD responses to the task were tested using an axiomatic approach to identify areas that may code pain PE adaptively.

**Results:** The left dorsal anterior insula demonstrated a pattern of response consistent with adaptively-coded pain PE. Signals from this area were sensitive to both predicted pain magnitudes on the instigation of expectation, and the unexpectedness of pain delivery. Crucially however, the response at pain delivery was consistent with the local context of the pain stimulation, rather than the absolute magnitude of delivered pain, a pattern suggestive of an adaptively-coded PE signal.

**Conclusions:** The study advances our understanding of the neural basis of pain prediction. Alongside existing evidence that the periaqueductal grey codes pain PE and the posterior insula codes pain magnitude, the results highlight a distinct contribution of the left dorsal anterior insula in the processing of pain.

**Significance statement:** Although there is behavioural evidence that pain is coded adaptively, the neural mechanisms serving this process are not well understood. This study used functional MRI to provide the first evidence that the left dorsal anterior insula, an area associated with aversive learning, responds to pain in a manner consistent with the adaptive coding of pain prediction error. This study aids our understanding of the neural basis of subjective pain representation, and thus can contribute to the advancement of analgesic treatments.

## Introduction

A major function of the nervous system is to learn the relationship between stimuli, and to identify, and respond accordingly, when expected relationships are confounded. According to the reinforcement learning framework (Sutton & Barto, 1998), prediction error (PE) is a key part of this adaptive process. PEs reflect the difference between the expected and delivered stimuli. A neural signal that corresponds to PE is generated when an expected stimulus is not delivered, and functions as a form of feedback, updating the cognitive schema that governs subsequent predictions about the world.

PE is typically computed according to reinforcement learning models (Sutton & Barto, 1998). A signal coding this “computational PE” should adhere to the following three axioms (Caplin & Dean, 2008; Rutledge et al., 2010):

Axiom 1: The signal should differentiate the magnitude of outcomes in a consistent order, independent of their probability (e.g. higher signal for outcome delivery vs outcome omission).
Axiom 2: The signal should reflect outcome likelihood consistently (e.g. lower signals for more expected outcomes).
Axiom 3: The signal should not differentiate between fully-expected outcomes.

Another, related function of the nervous system is to quickly identify the subjective value of stimuli. A problem with such a task is that the range of stimulus intensities is wide, while the processing range of neurons is limited. To reduce the computational demands of determining values, the brain is thought to engage in ‘adaptive coding’, where the sensitivity of some neurons to value is modulated by contextual information (Seymour & McClure, 2008). A neural signal that satisfies the computational PE axioms (listed above) independent of context, is sensitive to the absolute magnitude of the delivered stimuli. In contrast, an ‘adaptively-coded’ PE signal would produce a signal that is scaled to the range of magnitudes that could possibly be experienced in the local context, independent of the absolute magnitude of the delivered stimulus. Table 1 and Figure 1 illustrates how the adaptively-coded PE diverges from the computational PE.

**Figure 1.**
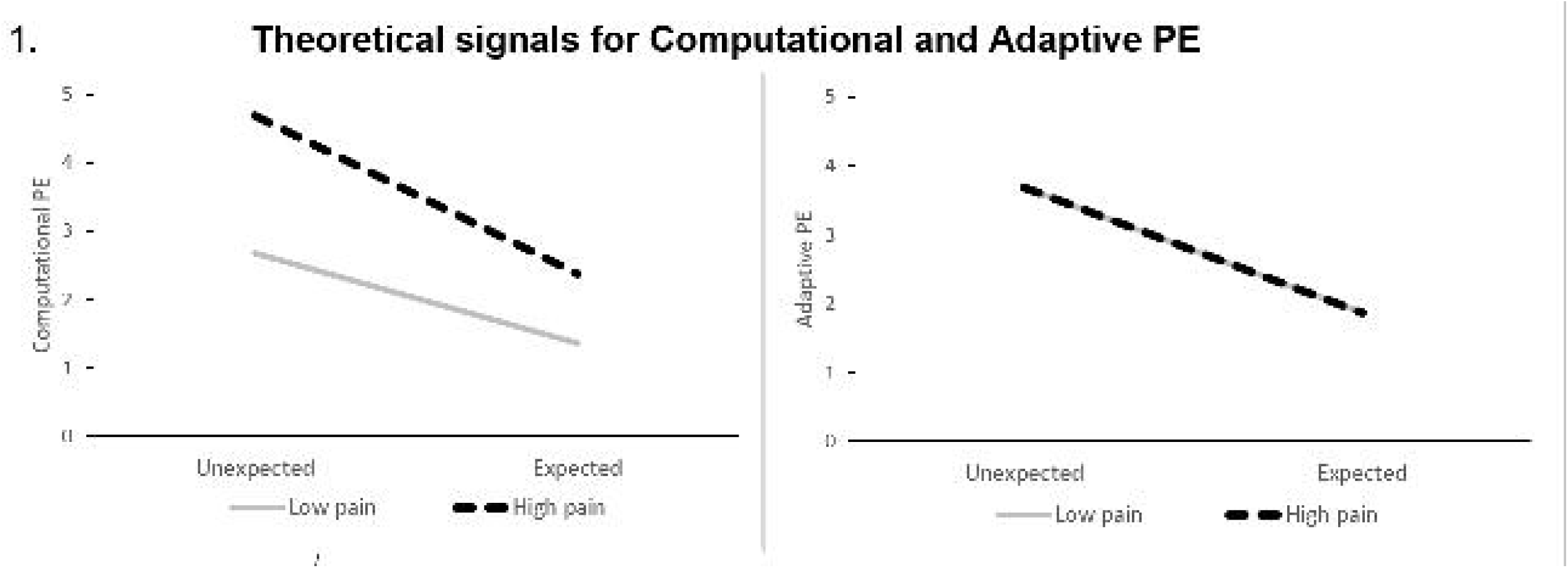
Theoretical levels for computational and adaptively-coded pain PE signals based on the current experimental design. The calculation of the values is given in Table 1. The left panel depicts the computational PE signal. All else held equal, computational PEs will be larger when pain is of high (vs low) magnitude and will also be larger when the pain is unexpected (vs expected). The right panel depicts the adaptively-coded PE signal. As with the computational PE, the adaptively-coded PE is higher when the pain is unexpected (vs expected). However, given the experimental design used here, the adaptively-coded PE will scale expectations from the cues such that either high or low pain are recoded as being the top of the pain magnitude range. Thus, a signal that expresses adaptively-coded PE will not be sensitive to pain magnitude in the current experiment.

**Table 1.**
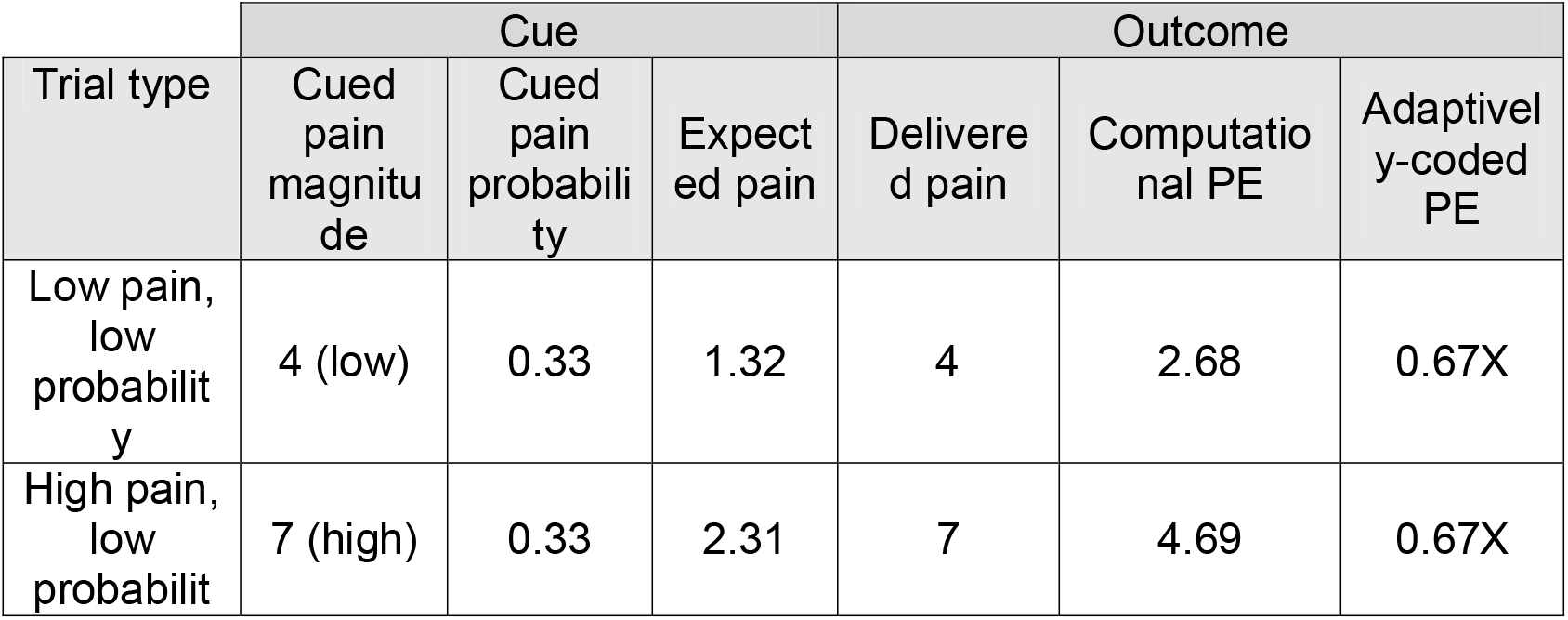
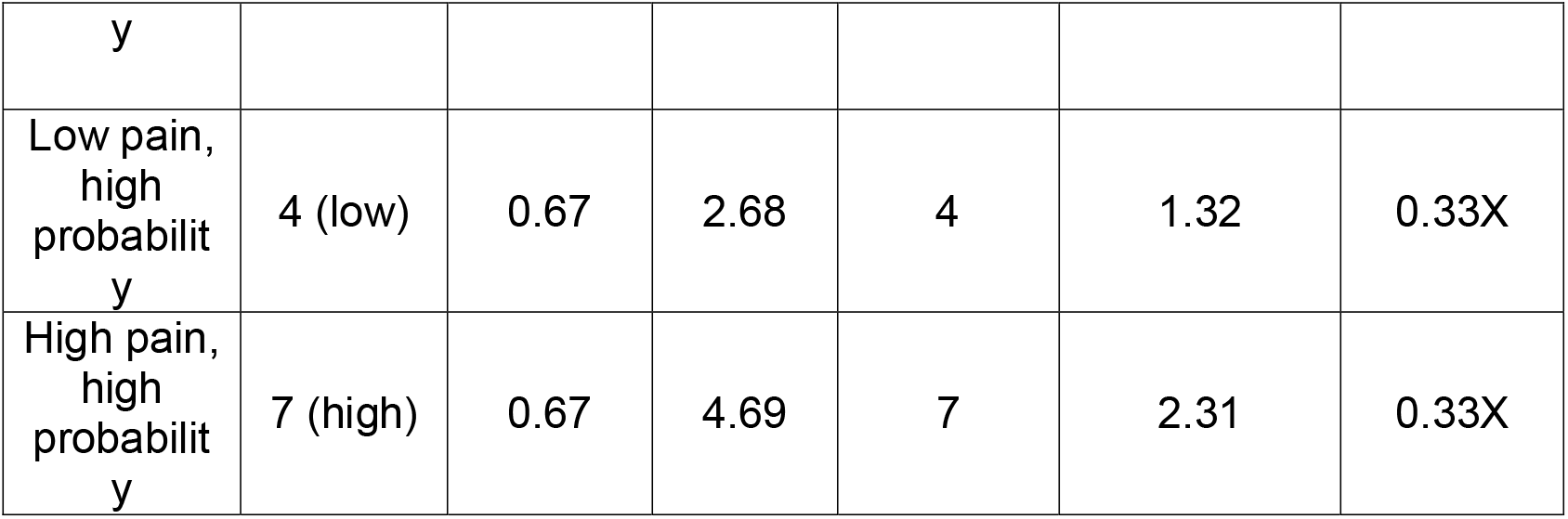
Example of the calculation of computational and adaptively-coded PE. shows the prediction error (PE) computation for trials where the stimulation is delivered. The pain magnitude was calibrated for each individual using a 1-10 scale where 4 was the pain threshold and 7 was the maximum pain tolerance level. Pain magnitude and pain probability were cued using visual symbols depicted in Figure 2. Expected pain was computed by multiplying magnitude and probability, according to the standard reinforcement learning procedure and following from expected utility theory. The computational PE was computed by subtracting expected pain from delivered pain magnitude. In the current experiment only one magnitude of pain could be delivered in each trial. This meant that, regardless of its magnitude, the pain stimulation was always the most adverse outcome possible within the context of each trial. Since the adaptively-coded PE is sensitive to the local context of the trial, the magnitude of the potential stimulation would not therefore impact the adaptively-coded PE in this paradigm. Instead, the adaptively-coded PE would be proportional to the probability of delivery for the sole pain level that was possible within the trial (computed as 1 – probability). This is multiplied by ‘X’, which signifies an unknown scalar representing the participant’s sensitivity to pain expectation. For the purposes of plotting the adaptively scaled PE in Figure 1, X has nominally been set to 5.5 (the average of the two pain levels delivered).

For rewarding stimuli, adaptive coding is implemented through dopaminergic neuronal firing (Tobler et al., 2005). Tobler and colleagues demonstrated adaptively-coded PE in Macaques. In each trial of their experiment Macaques observed one of three cues that signalled a 50% chance of receiving reward, with each cue signalling one reward magnitude only (low, medium, or large). Cue onsets triggered a midbrain dopaminergic signal proportional to the magnitude of the reward it predicted, namely, it scaled to the size of the potential reward. On trials where the reward was delivered, higher-than-baseline response in the same neurons reflected reward PE. Crucially for an adaptive signal, this response to the reward outcome no longer distinguished reward magnitudes, remaining the same regardless of actual reward delivered. This pattern of responses reflects an adaptively-coded PE, as in each individual trial the delivery of the reward was the best outcome (compared to nonreward), despite the reward magnitude being lower in some trails than in others.

In humans, evidence suggests that adaptively-coded reward prediction error is expressed in the ventral striatum (Park et al., 2012). As with reward, PEs triggered by the prediction or experience of aversive stimuli, such as pain, are also coded by dopamine neurons (Lammel et al., 2011; Schultz, 2016). Computational pain PE correlates with activation in the human ventral striatum and insula (Geuter et al., 2017; Seymour et al., 2004; Shih et al., 2019), although only in the periaqueductal grey does the signal adhere to all three axioms (Roy et al., 2014). As regards adaptive pain responses, there is ample evidence that local context influences behavioural responses to pain so that they scale with expectations (Atlas & Wager, 2012; Tracey, 2010). Behavioural evidence specific for adaptive coding of pain is evident in that participants pay more for relief of moderate pain when the pain intensities they could receive range from low to moderate, compared when they range from moderate to high (Vlaev et al., 2009; Winston et al., 2014). There is however, controversy about the neural mechanism that enables adaptive coding of pain, with research implicating either the insula and ACC (Leknes et al., 2013), or the orbitofrontal cortex (Winston et al., 2014); while one study reported null neural effects (Bauch et al., 2017). As these previous studies have not focused specifically on whether a region represents pain PEs in an adaptively-coded manner, the current study was designed to examine whether neural evidence for adaptive coding of pain PEs can be acquired by closely following Tobler et al.’s design. To this end a paradigm was implemented where four cues predicted either high or low pain intensity with either high or low probability. Crucially, as with the Tobler study, the cues presented at trial onset only predicted the delivery (or omission) of one intensity of stimulation. Thus, in the local context of each trial, the delivery of the pain stimulation was the worst outcome (compared to the alternative, pain omission), regardless of the intensity of the actual stimulation. It was hypothesized, based on Tobler et al., that a brain area that records PE would, at the point during which expectations are set, produce a response that scales with the intensity of the predicted stimulus. Given that we know that pain, especially when unexpected, increases the BOLD response, the activation of an area demonstrating an adaptively-coded pain PE would meet the following four requirements:

1. At outcome, respond more to delivered stimulations versus omitted stimulations (reflecting Axiom 1 of the Computational PE)
2. At outcome, respond more to unexpected (versus expected) stimulations (reflecting Axiom 2 of the Computational PE)
3. At outcome the effect of expectancy, namely, the greater response to the unexpected stimulation compared to the expected stimulation (requirement 2) does not vary with the actual intensity of the delivered stimulus. This reflects the requirement of an adaptively-coded PE signal to be sensitive to the local context (Figure 1).
4. At cue onset, respond more to the high pain cue than the low pain cue (reflecting Axiom 1 of the computational PE and following the findings of Tobler et al., where the area coding adaptively-scaled reward PE also responded to cued reward level).

Note that with the experimental design utilised, it was not possible to test Axiom 3 of the computational PE. As pain delivery was necessarily probabilistic, no stimulation was ever fully expected.

## Methodology

### Experimental design

The experiment employed a 2X2 event-related design with pain magnitude (high vs. low) and pain probability (high vs. low) as the factors. The task resembled one previously employed to detect neural signature of pain and reward PEs on the scalp (Talmi et al., 2013). Participants viewed chance cues that predicted electric stimulation with a 33% or 67% probability. In each trial it was only possible to receive a stimulation of a single magnitude, with the cue colour indicating which magnitude this could be (Figure 2). Across the experiment two different magnitudes were used: a ‘high’ level, equivalent to the most intense level of stimulation participants had previously indicated they were willing to tolerate, and a ‘low’ level, equivalent to a stimulation level that participants had indicated was “just painful”. An event-related design was used such that every block contained each of the 4 trials type (high and low probability and high and low pain). The trials were designed to permit the cue to adaptively scale the participants’ expectations, such that when the cue signalled a chance of low magnitude stimulation participants expected low (or no) pain, and when the cue signalled a chance of high stimulation they expected high (or no) pain.

**Figure 2.**
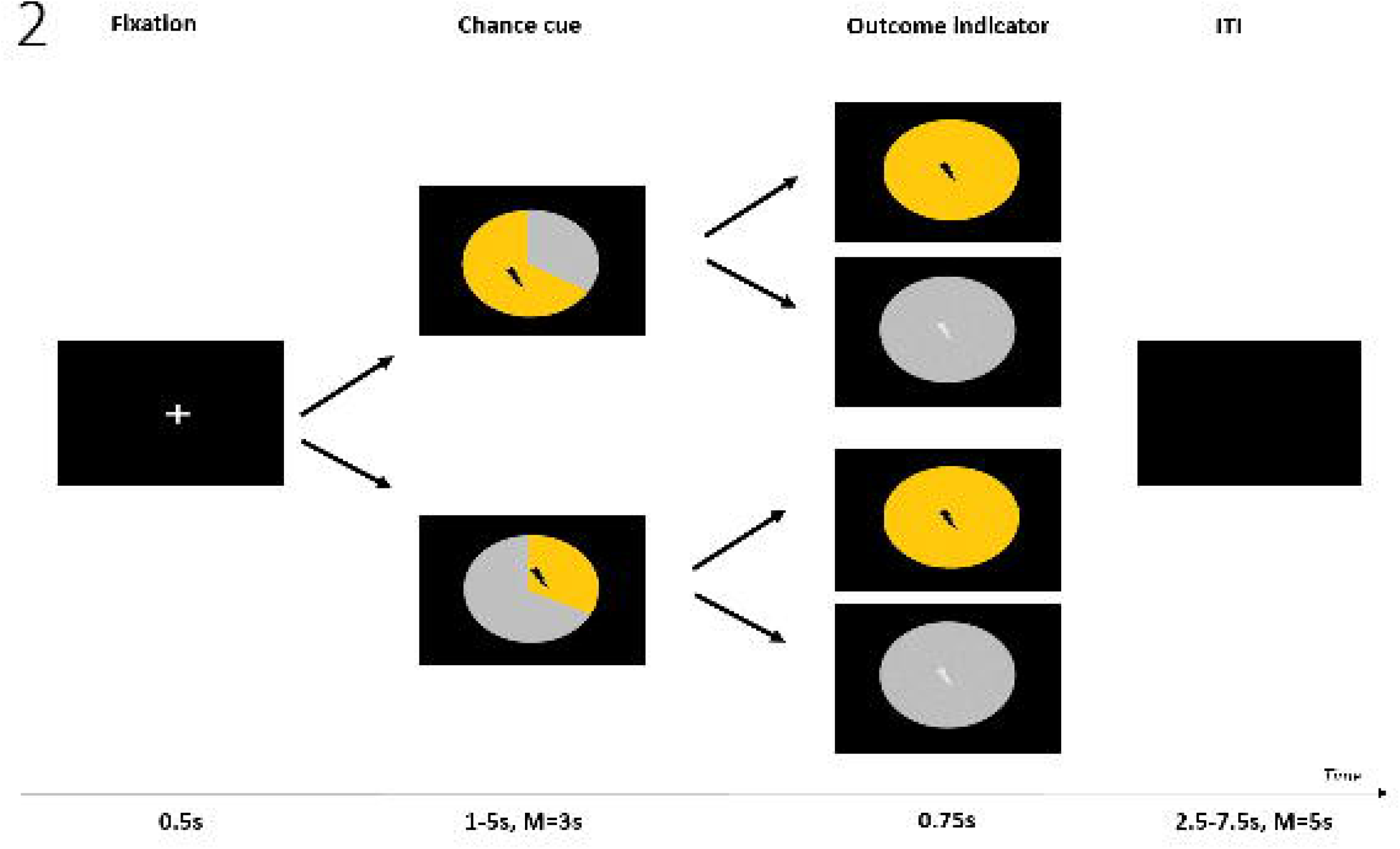
Graphical display of the timeline of a single trial. Participants saw a chance cue, where the non-grey portion signalled which pain magnitude could be delivered in the trial and with what probability (33% or 67%). Two non-grey colours (blue and yellow) were used to signal the high and low pain magnitudes. The figure depicts the cues one participant saw in the high pain condition (colours were counterbalanced across participants). Participants then saw an outcome indicator, where colours reflected whether the stimulation was, in fact, delivered. Fully coloured outcome indicators signalled pain delivery, while grey outcome indicators signalled pain omission.

### Participants

Twenty participants took part in the experiment, although data collection was aborted for one participant as they could not tolerate being in the scanner, leaving a final sample of 19 participants (11 female) between the ages of 18-36 (M=25.16, SD=4.35). This sample size was based on Hoskin et al. (2019) where a similar sample size was sufficient to reveal effects of expectations on pain experience, and with the sample sizes used by previous studies on adaptive pain PE (Bauch et al., 2017; Leknes et al., 2013; Winston et al., 2014). For two participants data was only collected for 3 (of the 4) blocks due to equipment failure. All participants were screened for any conditions that would prevent MR scanning, and for psychiatric and neurological history. Participants were proficient in English, had normal or corrected-to-normal hearing and vision, and did not take centrally-acting medication. The study received ethical approval from the University of Manchester ethics committee.

Further sample characteristics are described in Table 2.

**Table 2.**
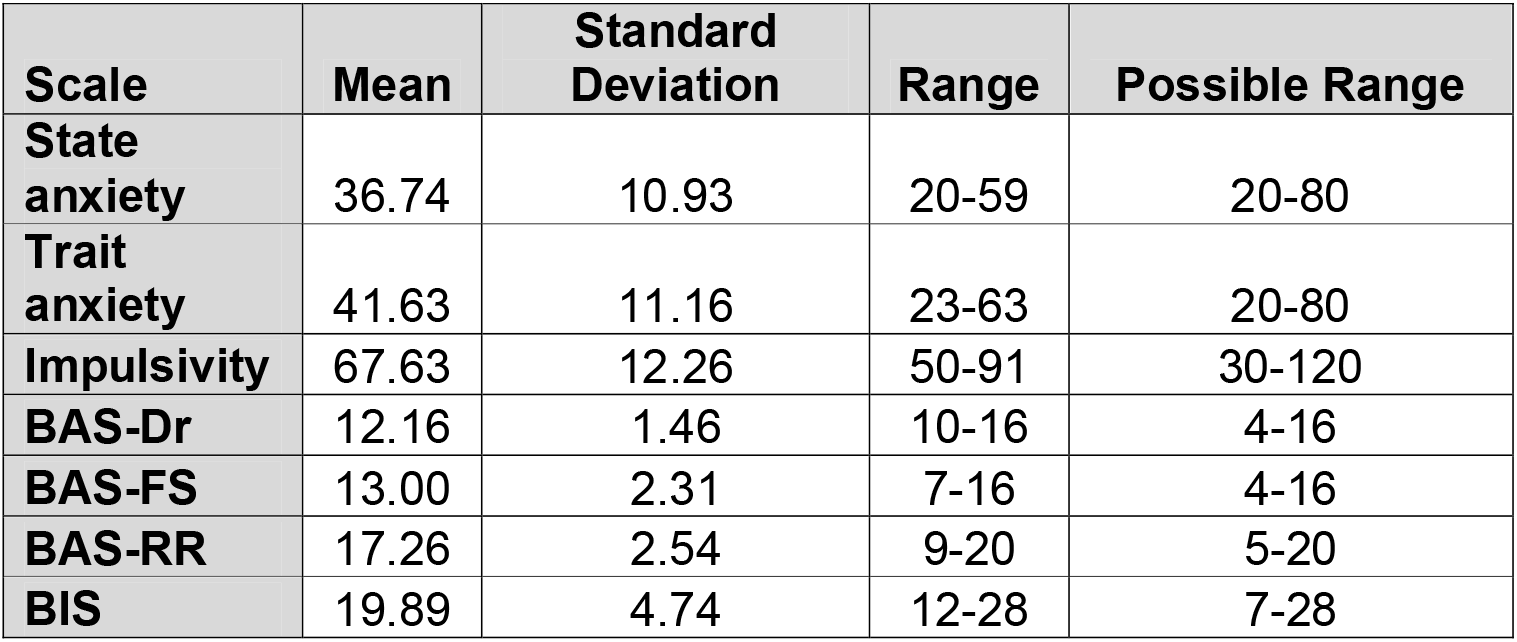
Sample characteristics (‘Range’ shows the lowest and highest actual scores, ‘Possible Range’ shows the lowest and highest possible scores).

| Scale | Mean | Standard Deviation | Range | Possible Range |
| --- | --- | --- | --- | --- |
| State anxiety | 36.74 | 10.93 | 20-59 | 20-80 |
| Trait anxiety | 41.63 | 11.16 | 23-63 | 20-80 |
| Impulsivity | 67.63 | 12.26 | 50-91 | 30-120 |
| BAS-Dr | 12.16 | 1.46 | 10-16 | 4-16 |
| BAS-FS | 13.00 | 2.31 | 7-16 | 4-16 |
| BAS-RR | 17.26 | 2.54 | 9-20 | 5-20 |
| BIS | 19.89 | 4.74 | 12-28 | 7-28 |

### Apparatus

#### Pain stimulation

The electrical stimulations were delivered to the back of the right hand via an in-house built ring electrode (Medical Physics, Salford Royal Hospital) attached to a Digitimer DS5 Isolated Bipolar Constant Current Stimulator (http://www.digitimer.com/). To counter the effect of the magnetic field of the MR scanner, the DS5 stimulator was placed within a custom-built Faraday cage (Medical Physics, Salford Royal Hospital). For reasons of participant safety, this stimulator was limited to delivering a maximum of 10mA during the experiment. To ensure adequate conductance between the electrode and the skin, the back of each participant’s hand was prepared with Nuprep Skin Preparation Gel and Ten20 Conductive Paste. The SCR electrodes were placed on the inside medial phalange of the second and fourth fingers of the participant’s left hand. The inputs to the DS5 machine were controlled via 1401plus data acquisition interface connected to a laptop running the Spike2 software (Cambridge Electronic Designs, Cambridge, UK). During the calibration procedure, the input signal from the 1401plus interface to the DS5 started from 0.2V and incremented at levels of 0.2V, up to a maximum of 5V. Using our set current range of 10mA, this driving signal was converted to current using the following equation: Input voltage/5V × 10mA = Output Current, so current levels experienced ranged from 0.4mA to 10mA. Both the laptop and the 1401plus machine were kept inside the scan control room during data collection. The 1401plus machine was connected to the Faraday cage using a BNC cable. To avoid interference between the Faraday cage and the scanner, the Faraday cage was kept a minimum of 2m from the scanner. The experiment was delivered via Cogent2000 on a Matlab platform (www.Mathworks.com).

#### Neuroimaging

Participants were scanned using a 3T Philips Achieva scanner, fitted with a Philips 32-channel receive-only coil. Whole-brain functional images were collected using a single-shot dual-echo protocol (Halai et al., 2014), with TR=3s, TE=12ms & 35ms, FOV=240,240,132, flip angle=85. In each volume, 33 slices of voxel size 3×3×4mm were collected in ascending order. Volumes were sampled at a 30-degree angle from the AC-PC line. Each functional scan included 145 whole brain volumes. Prior to the functional scans, a whole-brain T1-weighted anatomical scan was acquired from each participant (TR=8.4s TE=3.8s, flip angle=8).

### Procedure

Upon arrival participants read an information sheet and signed a consent form. Participants were then provided with comprehensive instructions outlining the task that was to be performed. Participants also filled out the following questionnaires (Table 2): Spielberger State-trait Anxiety Inventory (Spielberger et al., 1983), BIS and BAS scales (Carver & White, 1994) and the Barratt Impulsivity Scale (Patton et al., 1985) as part of a wider study investigating the relationship between anxiety, impulsivity and the neural response to pain. Results relating to this questionnaire data are not therefore discussed further here.

Prior to engaging in the main task, participants first completed a calibration procedure which was designed to identify the two levels of electrical stimulation to be used during the main task. An established calibration process (e.g. Brown et al., 2014; Hird et al., 2018) was adopted, during which participants received a succession of 5ms square wave electric stimulations. These stimulations started at a very low, barely perceptible level, and the current level increased very gradually (by incrementing the driving signal by 0.2V at each time, see ‘Apparatus’ section). Participants rated each stimulation on a scale from 0 – 10 where a score of 0 reflected not being able to feel the stimulation, 4 reflected a stimulation that was on the threshold of being painful, 7 related to a stimulation that was deemed ‘painful but still tolerable’ and 10 related to ‘unbearable pain’. The procedure was terminated once the participant reported the level of pain as being equivalent to ‘7’ on the scale. The procedure was performed twice, both times with a one-step-up method, to allow for initial habituation/sensitisation to the stimulation. The voltage levels rated as ‘4’ and ‘7’ on the second scaling were used for the ‘low’ and ‘high’ pain stimulation magnitude levels during the main task. The calibration procedure ensured that the pain levels were psychologically equivalent across participants.

Participants first completed 1 block of 60 trails outside the scanner to ensure that they understood the task. Inside the scanner participants completed a further 4 blocks of 60 trials. Each block contained 15 trials of each of the 4 cues. The trial structure is depicted in Figure 2. Each trial began with a 0.5s fixation cross before a ‘chance’ cue appeared in the form of a two-segment pie chart, with one segment always being coloured grey. Each chance cue signalled two attributes of the outcome of the trial. The magnitude of the pain that could be administered in the trial was signalled through the colour of the non-grey segment, with (across the experiment) one colour signalling high pain and the other low pain (colours used were blue and yellow and their assignment to high and low were counterbalanced across participants). The grey segment was used to signal the possibility of receiving no stimulation. The probability of receiving pain in each trial was therefore signalled through the portion size of the coloured (non-grey) segment. A chart where 67% was coloured signalled a 67% chance of receiving the pain stimulation in the trial (high probability); whereas a chart where the 33% portion was coloured signalled a 33% chance of receiving the pain stimulation (low probability). The cues accurately signalled the probability of the stimulation being delivered both across the entire experiment and within each block (e.g. out of the 15 presentations of each 33% cue in a block, 5 would results in stimulations and 10 in no stimulations). While the delivery or omission of the outcome was probabilistic, the chance cue clarified that only an outcome of a single magnitude (high in half the trials, low in the other half) was possible, with the only other possible outcome being the omission of the stimulation. Thus, a crucial feature of the task was that participants never needed to consider a possibility that they may get either high or low pain. By setting participants’ expectations in this way, the chance cue allowed the pain PE to be adaptively coded.

After the cue, an interval of between 1 and 5s occurred before an ‘outcome indicator’ appeared, taking the form of a one-segment pie chart, coloured according to the outcome of the trial. In trials where stimulation was omitted, the outcome indicator was grey. In trials where stimulation was delivered, the outcome indicator was coloured according to the magnitude delivered, with the indicator appearing at the onset of the stimulation. Outcome indicators were included in all trials to provide certainty as to the outcome for trials in which no stimulation was delivered, thus avoiding introducing confounds around temporal uncertainty. A jittered inter-trial interval of between 2.5s and 7.5s then occurred before the start of the next trial.

To ensure participants paid attention to the contingencies presented by the cues, they were asked to monitor the outcomes that followed each cue. At the end of each block participants were shown each chance cue again (order randomised) and asked whether they thought the cue accurately predicted the occurrence of the stimulations (possible responses: “yes” or “no”). Prior to starting the task, participants were informed of the nature of the chance cues and outcome indicators. To ensure they engaged with the monitoring task, participants were not informed that the probabilities represented on the chance cues were in fact accurate.

### Analysis

#### Pre-processing

MATLAB and SPM12 were used to pre-process the raw scans and complete analysis of the resulting data in the 1^st^ and 2^nd^ level GLM (general linear model). Raw PARREC files of the structural and functional MRI data were converted into file formats suitable for SPM12 (.img/.hdr & .nii files) in order to be preprocessed using MATLAB code. A standard neuroimaging pipeline using a mass-univariate approach was subsequently implemented on these data-files/scans. Realignment/motion-correction was applied first, followed by slice timing correction using the central slice for reference. Spatial normalisation was then applied, with the functional data being normalised into a standard stereotactic MNI (Montreal Neurological Institute) space resampled to 3 × 3 × 4mm^3^ voxels before being spatially smoothed using an 8-mm [8 8 8] full-width at half-maximum (FWHM) Gaussian kernel in order to optimise sensitivity (Ashburner et al., 2016). Following pre-processing the resulting data-files were specified in the 1st level GLM. A 128s high-pass filter was used to reduce the effects of MRI scanner drift. Inclusion of temporal derivatives in subject’s 1st-level analysis were considered, however, this was ultimately deemed counter-productive due to evidence (Della-Maggiore et al., 2002; Sladky et al., 2011) that including temporal derivatives in paradigms centred on response latencies >1s can diminish power, thereby directly subverting the increase in sensitivity provided by slice-timing correction. The literature dedicated to onset latency and delays of the hemodynamic response function (hrf) in response to pain stimuli are quite limited (Cauda et al., 2014; Pomares et al., 2013) with only Cauda et al. reporting that the canonical hrf may not be well-suited to capturing the BOLD response of measured mechanical pain. As this paradigm used transcutaneous electric nerve stimulation, not mechanical pain, no derivatives were included in the GLM, and the canonical hrf was used, as in the majority of published research on pain.

#### Individual models

Individual GLMs were constructed for each participant, encompassing all completed runs. Each run was modelled with 9 regressors. Four regressors corresponded to chance cues, crossing the factors pain magnitude (high vs. low) and pain probability (high vs. low) and another 4 regressors corresponded to pain outcomes (i.e. the delivery of a stimulation), crossing the same factors. Note that when the chance cue indicated that pain probability was high, the subsequent delivery of pain was taken to correspond to ‘expected pain’, while when the chance cue indicated that the pain probability was low, the subsequent delivery of pain was taken to correspond to ‘unexpected pain’. The 9^th^ regressor modelled all non-pain outcomes (i.e. the presentation of the outcome indicator that signalled that no pain would be delivered), following from previous work where non-pain outcomes did not trigger a pain PE (Geuter et al., 2017). Six motion parameters were also included for each run.

#### Group analysis

Two second-level ANOVA tests were constructed. The *‘pain anticipation’* model analysed the response to chance cues and the *‘pain outcome’* model analysed the response to pain outcomes, each crossing the factors pain magnitude x pain probability. Main effects were examined using *t*-contrasts and interactions using f-contrasts, with *p*<.05 for initial voxel selection, and *FWE* <.05 to define statistically-significant voxels. The search volume for all group analyses was constrained to Regions Of Interest (ROIs) that were sensitive to pain. These ROIs were defined functionally through a one-sample *t*-test, which contrasted the response to pain outcomes (an average across all four pain outcome regressors) to responses to non-pain outcomes (the 9^th^ regressor). A conservative threshold of *FWE*<.05 was used to define the functional ROI mask. The mask was then used to constrain the search volume in all reported group analyses, unless otherwise stated.

## Results

The response to the monitoring question at the end of the block showed that participants were more likely to say they thought the cue accurately predicted the probability of pain. Across all cues, 64% of the responses to the monitoring question were “yes”, with the majority of responses also being “yes” to each cue individually.

To identify areas that adhered to requirement 1 of the adaptive PE (and Axiom 1 of the computational PE) the ‘pain outcome’ model was used to identify regions that were sensitive to pain outcomes compared to non-pain outcomes. This contrast revealed that the anterior and posterior left insula, anterior right insula, middle and anterior cingulate cortex, as well as activations in the supramarginal gyrus, angular gyrus, supplementary motor cortex, and inferior frontal gyrus met this requirement. For completion, the opposite contrast (non-pain outcomes > pain outcomes) was also examined, a contrast which could involve psychological relief. This analysis did not identify any statistically significant voxels.

As a manipulation check, we used the *‘pain outcome’* model to examine the main effect of pain magnitude (high vs low). The analysis identified peaks in the posterior insula bilaterally (Figure 3) corresponding to greater activation to high vs low pain. No areas showed greater activation for the delivery of the low pain stimulus.

**Figure 3.**
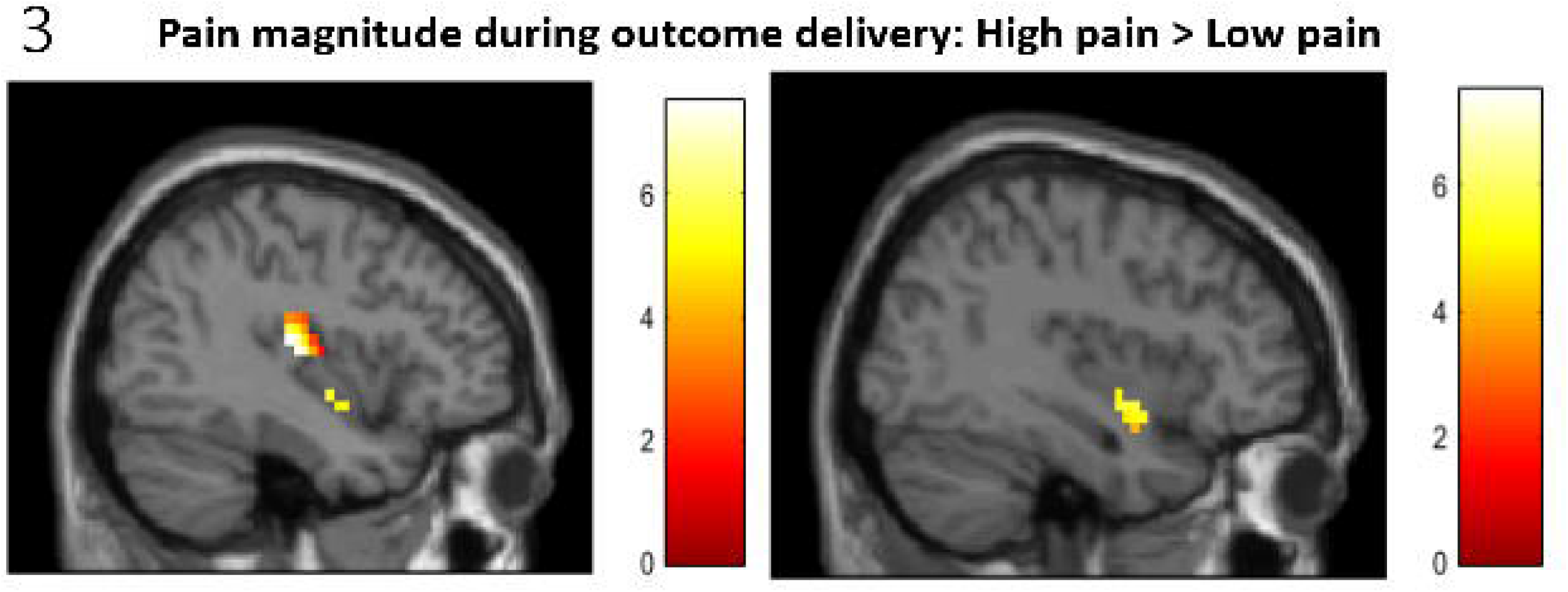
Response to pain magnitude, high > low pain. The sagittal slices (right panel: X=−39, left panel: X=39) show that in the bilateral posterior insula, signal was larger when the pain delivered was of high magnitude, compared to when it was low, *FWE<.05.*

Regarding the main analysis, figures 4A and 4B depict the key main effects: the effect of pain magnitude at anticipation, and the effect of pain probability at outcome (in a 2*2 factorial design, ‘main effects’ refer to the effect of one factor, collapsing across the levels of the other factor). Figure 4C shows the parameter estimates for these main effects at both anticipation and at outcome for a peak insula voxel. Figure 4D depicts the key simple effects, which are discussed in more detail below (in a 2*2 factorial design, ‘simple effects’ refer to the effect of one factor within one level of the other factor).

**Figure 4.**
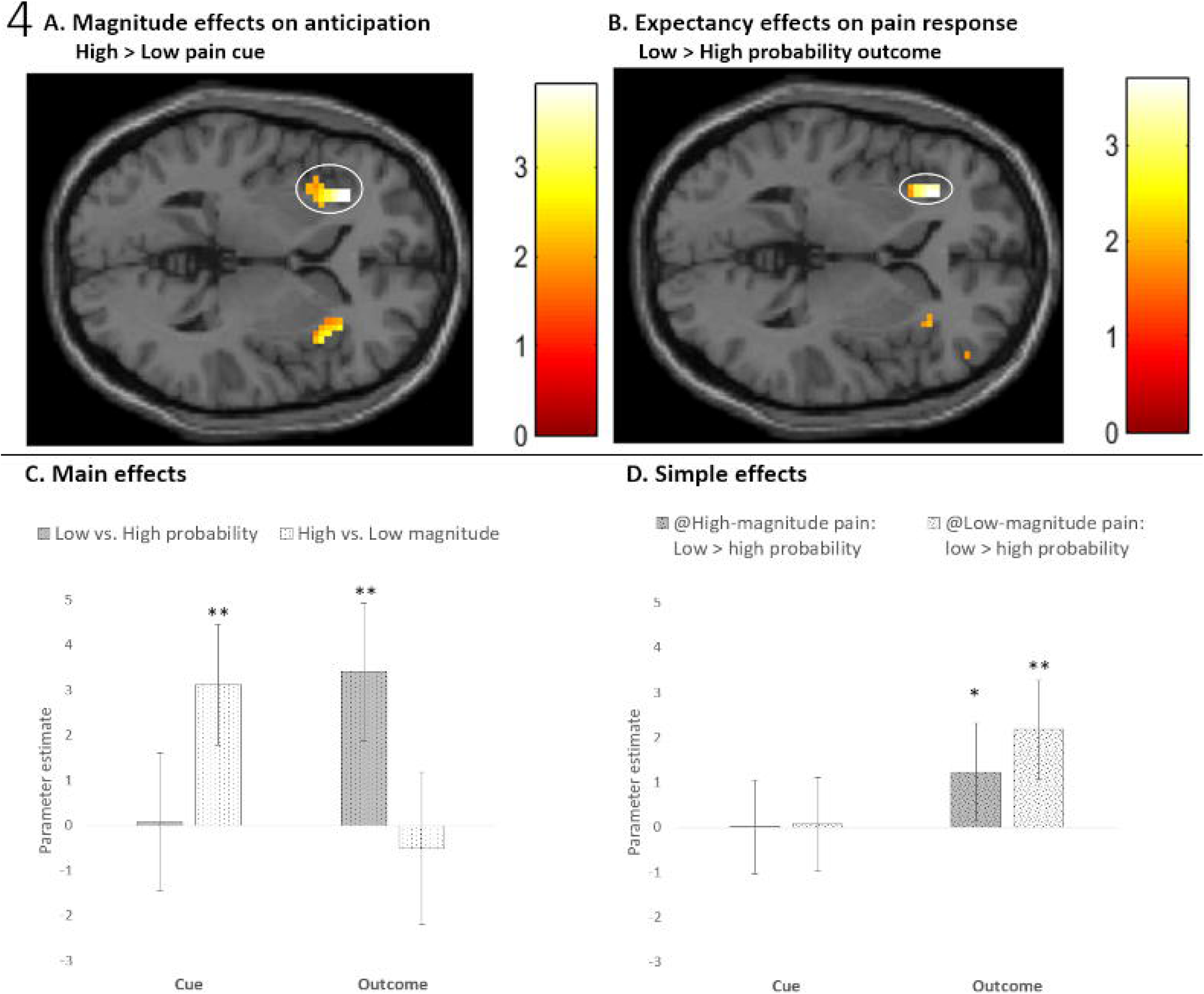
Effects of pain probability and magnitude on pain anticipation and pain outcome in the left anterior insula. A. Effect of pain magnitude on pain anticipation, with those clusters surviving correction for multiple comparison (FWE<.05) circled. Analysis of the pain anticipation model shows that signal in the left dorsal anterior insula (X=−30, Y=29, Z = 2) was higher when the pain cue signalled high pain, compared to when it signalled low pain. A. Effect of pain expectancy on pain outcome, with those clusters surviving correction for multiple comparison (FWE<.05) circled. Analysis of the pain outcome model shows that signal in left dorsal anterior insula (X=−30, Y=26, Z =2) was higher when delivered pain was unexpected (33% likely), compared to when it was expected (66% likely). B. Parameter estimates of the main effects of pain probability (dark bars) and pain magnitude (light bars) in response to the cue (anticipation: left) and pain delivery (outcome: right) in the left dorsal anterior insula (X=−30, Y=26, Z =2), FWE<.05. This region was responsive to the magnitude of pain when expectations were set at the time the chance cue was presented (high pain cue > low pain cue), but not to the magnitude of experienced pain at outcome (null main effect of pain magnitude at outcome). During pain delivery the same region was responsive to pain probability (unexpected pain > expected pain). Error bars represent the 90% confidence interval. **C.** Simple effects of pain probability at cue (anticipation: left) and outcome (right) in the left insula (X=−30, Y=26, Z =2) when the pain magnitude was high (dark bars) or low (light bars). Error bars represent the 90% confidence interval. *p<.05, ** FWE<.05 in a paired t-test (see text).

To test requirement 2 of the adaptive PE (Axiom 2 of the computational model), we examined the main effect of pain probability on the response to the delivery of pain, using the *‘pain outcome’* model. The analysis identified a single significant peak, in the left dorsal anterior insula (Chang et al., 2013), where response to unexpected pain was greater than response to expected pain (Table 3 and Figure 4B). The effect of probability at outcome was in the same direction (low probability > high probability) for each pain magnitude, and that the 90% confidence interval excluded zero for both (Figure 4D). We verified this result by constructing two new flexible factorial models, one for each pain magnitude (high or low), and each with two vectors, one for high and one for low probability. Analysis was limited to the significant left dorsal anterior insula cluster. Paired t-tests suggested that the probability effect was significant there for high pain (peak X=−30, Y=23, Z=2) at an uncorrected p<.05 (FWE<.1), and for low pain (peak X=−30, Y=29, Z=6) at FWE<.05. Taken together these results suggest that this region adheres to both Axioms 1 and 2 of the computational model of PE and requirements 1 and 2 of the adaptively-coded PE. No area responded more to expected pain than unexpected pain.

**Table 3.**
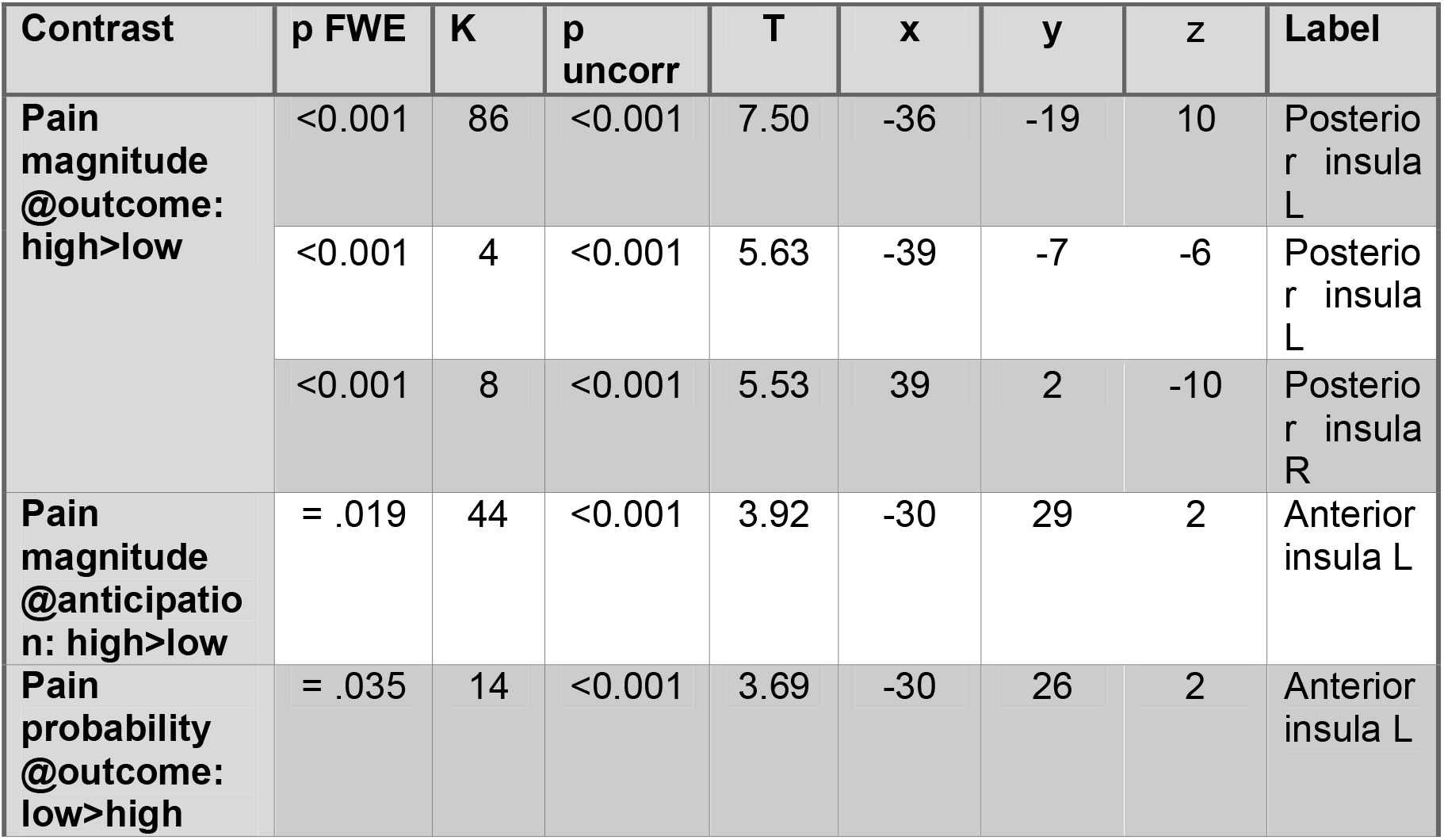
Differences in BOLD signal change found in the main effects from the ANOVAs for the ‘anticipation’ and ‘outcome’ models. The table shows differences in BOLD signal change found in the main effects from the ANOVAs for the ‘anticipation’ and ‘outcome’ models, inclusively masked by the contrast pain delivered>pain omitted. Both models involved the factors of pain magnitude (high, low) and probability (high, low). Only activations that survived the family-wise error (FWE) threshold of <=.05 are included. We entered as dependent variables the T map for each subject, either during the outcome delivery or the chance cue, averaged across sessions. The activations relating to the first contrast (pain magnitude @ outcome) are also shown in Figure 3, while the activations in the second and third contrasts are shown in Figures 4a and b respectively. Note that a more lenient threshold was used for voxel detection in the figures, in comparison to the strict FWE threshold, which corrects for multiple comparisons, used in this table. Clusters therefore appear in the figure which do not appear in this table.

| Contrast | p FWE | K | p uncorr | T | x | y | z | Label |
| --- | --- | --- | --- | --- | --- | --- | --- | --- |
| **Pain magnitude @outcome: high>low** | <0.001 | 86 | <0.001 | 7.50 | -36 | -19 | 10 | Posterior insula L |
|  | <0.001 | 4 | <0.001 | 5.63 | -39 | -7 | -6 | Posterior insula L |
|  | <0.001 | 8 | <0.001 | 5.53 | 39 | 2 | -10 | Posterior insula R |
| **Pain magnitude @anticipation: high>low** | = .019 | 44 | <0.001 | 3.92 | -30 | 29 | 2 | Anterior insula L |
| **Pain probability @outcome: low>high** | = .035 | 14 | <0.001 | 3.69 | -30 | 26 | 2 | Anterior insula L |

To test requirement 3 of the adaptive PE, the interaction between probability and magnitude within the pain outcome model was assessed. No areas exhibited a statistically significant interaction effect at outcome between pain magnitude and probability, thus suggesting that the left dorsal anterior insula also satisfies requirement 3 of the adaptively-coded PE. A signal that corresponds to the computational PE pain signal should take the form of stronger response to unexpected pain when it is high magnitude, compared to when it is low (Figure 1). Figure 4D shows that in the left dorsal anterior insula peak, the pain magnitude contrast of the PE at outcome was (numerically) in the opposite direction. This suggests that the failure to find an interaction between the effects of pain magnitude and probability at outcome was unlikely to be due to a lack of power.

Finally, to test requirement 4 we examined the effects of pain probability and pain magnitude on the response to chance cues (Table 3), using the *‘pain anticipation’* model. The only significant response to the main effect of pain magnitude was again observed in the left dorsal anterior insula, where activation was stronger for chance cues that signalled high pain, compared to those that signalled low pain (Figure 4A and 4C). Thus, this area also satisfied requirement 4 of the adaptively-coded PE. Neither the main effect of pain probability nor the interaction influenced pain anticipation significantly within the functional ROI mask.

To summarise the findings in the dorsal anterior insula, Figure 4C illustrates that the pain magnitude contrast (light-shaded columns) was significant only during anticipation, and that the pain expectancy contrast (dark-shaded columns) was only significant during outcome (pain delivery). Figure 4D illustrates that the simple effect of pain expectancy at outcome was significant and similar in magnitude both when pain was high and when pain was low. This pattern suggests that a region in the dorsal anterior insula expresses adaptively-coded pain PE.

For completion, we examined whether pain magnitude or probability in response to chance cues, or the pain outcomes, activated any additional regions outside of the functional ROI mask. For this purpose, we used a conservative *FWE*<.05 for whole-brain search. Anticipating high (vs low) pain activated the right hippocampus and the right inferior frontal gyrus. The whole-brain analysis of pain outcomes did not uncover any additional activations.

## Discussion

Signal in the left dorsal anterior insula represented adaptively-coded prediction error (PE) of pain, defined according to an axiomatic approach such that it should satisfy the 4 requirements listed in the Introduction. As expected from a region sensitive to pain PE, upon cue presentation, the left dorsal anterior insula responded more strongly to the anticipation of high rather than low pain (requirement 4). Additionally, and again as to be expected from a region sensitive to pain PE, upon pain delivery, the signal in this region was stronger for low-probability (i.e. unexpected) pain compared to high-probability pain (requirement 2). Crucially, despite its sensitivity to pain magnitude when it was predicted, and pain probability when it was actually delivered, this region was insensitive to the magnitude of pain during delivery (requirement 3). The pain response signal was similar when either low or high pain were delivered, but significantly greater when the pain was unexpected. This result suggests that the chance cue scaled the subsequent response to pain PE, such that the dorsal anterior insula responded similarly to the highest-possible pain in the local context of the trial. Taken together, this pattern corresponds to the pattern reported by Tobler et al. (2005) which established adaptive scaling of prediction error for reward. Here we show, for the first time, a similar pattern for pain PEs.

While the dorsal anterior insula signalled adaptively-coded pain PEs, the posterior insula was sensitive to delivered pain intensity. Together, both results complement elegant results reported by Geuter and colleagues, where the signal in the anterior insula reflected the sum of cued pain expectations and pain PEs, while the posterior insula and parietal operculum coded for pain stimulation intensity (Geuter et al., 2017). PE is a signed quantity; either reward or pain delivery outcomes are considered to cause positive PE, and either omitted reward or omitted pain are considered to cause negative PE. In both our study and that of Geuter and colleagues, the focus was on positively-signed pain PE. Shih et al (2019) compared negative and positive PE for aversive stimuli and presented findings that suggest separate neural substrates for each. They found no areas that coded both negative and positive PE, but the anterior insula exhibited a higher BOLD response for positive PE, as in our study, and the anterior cingulate for negative PE. Interestingly, although they did not observe regions that expressed both positive and negative computational PE signal, Shih et al. found that the connectivity of both the insula and the ACC with the PAG increased with respect to the PE regardless of its sign. This result aligns nicely with Roy et al.’s findings that the PAG expressed the computational pain PE, which corresponds to computational reinforcement learning models where it is insensitive to local context.

This study advances understanding of the neural mechanism that serves the adaptive coding of pain, which is less comprehensive than our understanding of adaptive coding of reward (Vlaev et al., 2009). Despite evidence that reward is coded adaptively, and for shared neural mechanism for adaptively coding monetary gain and loss (Nieuwenhuis et al., 2005), it is, in principle, possible that the brain does not need to represent pain in an adaptively-coded manner and can instead represent all biologically-feasible levels of pain, either because they span a more limited range than all possible rewards, or because the evolutionary significance of pain, compared to reward, has caused the brain to represent pain more accurately. Nevertheless, adaptive coding of pain reflects an influence of pain expectations, and there is ample evidence that pain perception is influenced by expectations (Atlas & Wager, 2012; Tracey, 2010), that expectations are clinically relevant (Buchel et al., 2014), that they may exert a stronger impact on pain perception than the noxious stimulation itself (Lim et al., 2020) and that many of its detailed characteristics have unique influence on the experience of pain (Hoskin et al., 2019; Watkinson et al., 2013). An understanding of the neural mechanisms of adaptively-coded pain PE is important as it can help us harness the cognitive system to decrease pain, as in cognitive therapy for chronic pain.

There is also elegant behavioural evidence for adaptive coding of pain, reviewed in the introduction (Vlaev et al., 2009; Winston et al., 2014). The first study to examine the neural mechanism of adaptive coding of pain was conducted by Leknes et al. (2013). They presented participants with two cues, one that predicted a 50% probability of either high or moderate thermal pain, and one that predicted a 50% probability of either low or moderate pain. A comparison of the response to the moderate pain stimulation when it followed each cue revealed that activity in the insula and anterior cingulate cortex (ACC) was lower when the moderate stimulation had been preceded by the cue predicting high pain, suggesting that these regions represented adaptively-coded pain response. However, Leknes et al presented the two cues in separate blocks, contrasting the response to the moderate pain across the blocks. It is not therefore clear to what extent the activation found might reflect a generalised response to the block context, rather than a PE signal specific to the presentation of the moderate stimuli. Using a similar procedure, Winston et al (2014) found that activity in the lateral OFC, but not the Insula or ACC, reflected an adaptively-coded pain response. Although each individual trial in the Winston study involved a single pain level, individual blocks only involved two levels of stimuli (to create the local context for the PE), again potentially introducing ‘block effects’ to the contrasts. Interestingly, when Bauch and colleagues utilised an event-related variant of Leknes (2013) block design, they were unable to find any area which produced a significant adaptively-coded pain response. Here we used a design that was closer to that employed by Tobler et al. (2005), with a specific focus on adaptive coding of pain prediction errors, rather than on the scaling of pain value by expectations more broadly. This perhaps accounts for why our results differ from those of Winston and colleagues. Importantly an event-related methodology was used in the current study, meaning that confidence can be taken that the results do not reflect generalisation effects. The human neuroimaging literature reports a number of brain regions that correlate with computational pain PE, computed according to reinforcement learning definitions (Sutton & Barto, 2015), including the ventral striatum, anterior insula, and the cingulate cortex (Geuter et al., 2017; Lim et al., 2020; Roy et al., 2014; Seymour et al., 2004; Shih et al., 2019). These studies have not used an axiomatic approach (Caplin & Dean, 2008), so it is not known whether signal in these regions corresponds with all three axioms (Roy et al., 2014), nor whether it expresses quantities that are correlated with PE, such as the expected value of pain, shown to activate the ventral striatum and involve the anterior insula (Brooks et al., 2010; Palminteri et al., 2012). As in the current study, Geuter et al found activation consistent with Pain PE signals in the anterior insula, using heat rather than electrical pain stimulations. In that study the PE could not be adaptively coded because participants were informed that they could receive the least favourable, highest stimulation intensity in each trial, with the result that in the local context of each trial was always the same. This was avoided in the current study because while the cue signalled that the high stimulation intensity was the least favoured option in half the trials, in the other half, it signalled that the low pain stimulation was the least favoured option. The cue therefore created a local context, which ensured the ensuing PE signal could be adaptive scaled. Our results suggest that the PE found in the anterior insula by Geuter et al, may be subject to adaptive scaling.

The left anterior insula has been long thought to be important for the representation of learning. It has been found to express levels of deviation from expectation (Fouragnan et al., 2018), and correlate with PE for motivationally neutral perceptual stimuli (Nazimek et al., 2013). In terms of aversive learning, Palminteri and colleagues (Palminteri et al., 2012) were able to establish, using patients with lesions in the anterior insula, that the area is involved in updating the value of loss-predicting cues. The anterior insula is also associated with signalling the behavioural relevance of information, for example providing early responses to facial stimuli according to the relevance of the emotion expressed upon it (Frot et al., 2022). Since unexpected stimuli are likely to be more behaviourally relevant than expected ones, there is a natural overlap between signalling PE and behavioural relevance. Nevertheless, in the current study the response of the anterior insula was no different when unexpected pain was of low compared to high magnitude. As higher pain is more behaviourally relevant, this suggests that the results of the current study cannot solely be attributed to the insula’s function in signalling behavioural relevance.

One potential shortcoming of the experimental design is that it did not include a behavioural task that would allow responses to be recorded on each trial. This prevented an assessment of the behavioural impact of PE. Trial-by-trial pain ratings were not requested firstly because they were not necessary for the research question, and secondly to ensure that cognitive processes associated with the response to questions did not interact with the neural responses to outcomes. The prediction error signal is a computational quantity and its relevance to brain function is evidenced in a large number of electrophysiological and imaging studies (Rutledge et al., 2010; Schultz, 2016). Although PEs are essential for learning, whether they have any immediate behavioural impact is unclear. Indeed recent research has demonstrated that it is pain expectations, rather than PE, that influence pain perception during probabilistic processing (Nickel et al., 2022). Nevertheless, future studies may benefit from collection of further behavioural data, which could be useful for the purpose of manipulation check. For example, trial-by-trial pain ratings would help assess whether expectations that are purely focused on the likelihood of pain influence its experienced intensity.

At the end of each block participants were required to signal whether they thought the probabilities reflected on the chance cues corresponded to the frequency of pain that was actually delivered. This task was included to ensure participants paid attention to the cue-outcome relationships, and thus engaged with the probabilistic nature of the paradigm. Only a modest majority of these reports confirmed belief that the cues represented outcome likelihood accurately.

Participants may have responded in this way because they believed that the frequency of outcome did not conform exactly with the probabilities shown, even though in reality, it did. Future studies would benefit from using a more focussed task, for example asking participants to estimate the frequency with which each cue was followed by a stimulation. It would be interesting to see whether participants’ responses reveal systematic bias, for example a belief that the frequency of unexpected pain was higher than 33%.

Future research could also utilise a wider variety of probabilities. The current research only involved two levels of probability, High and Low. Although this was sufficient to test requirements 2 and 3 of the adaptively-coded PE, the use of more probability levels may increase the sensitivity of these tests. The use of more probability levels may also allow an assessment of the exact mathematical relationship between probability and PE and what difference in probability is required to produce a significant difference in PE.

A shortcoming of the experimental design was the use of the outcome period for the no-pain conditions as a baseline for the identification of the pain-sensitive regions. Although this contrast satisfies the first axiom of the computational PE, it is potentially contaminated with relief that might have been experienced on the trials where no stimulation was delivered. However, the opposite contrast (no pain >pain) did not reveal any significant results, suggesting that relief was not strongly experienced in the regions of interest. Furthermore, the other contrasts performed in the analyses were not associated with relief, such as the main effects of pain probability or magnitude. Therefore, we do not believe this shortcoming impacts the results presented. The region identified here as important for adaptive pain PE, the anterior insula, is widely considered to be involved in pain processing (Garcia-Larrea & Peyron, 2013).

In summary the results of the current research suggest that while the left anterior insula is sensitive to many aspects of aversive learning, and to quantities associated with the computational PE signal for pain, it may better represent an adaptively-coded pain prediction error.

## Acknowledgements

We thank Daniel Wilde for help with data analysis, and E. J. Hird for helpful comments.

## Author contribution

DT and RH designed the study. RH collected the data. DT and RH wrote the manuscript. All authors discussed the results and commented on the manuscript.

